# A Neural Speech Decoding Framework Leveraging Deep Learning and Speech Synthesis

**DOI:** 10.1101/2023.09.16.558028

**Authors:** Xupeng Chen, Ran Wang, Amirhossein Khalilian-Gourtani, Leyao Yu, Patricia Dugan, Daniel Friedman, Werner Doyle, Orrin Devinsky, Yao Wang, Adeen Flinker

**Affiliations:** Electrical and Computer Engineering Department, New York University, 370 Jay Street, Brooklyn, 11201, USA, Country; Neurology Department, New York University, 223 East 34th Street, Manhattan, 10016, USA, Country; Biomedical Engineering Department, New York University, 370 Jay Street, Brooklyn, 11201, USA, Country; Neurosurgery Department, New York University, 550 1st Avenue, Manhattan, 10016, USA, Country

**Keywords:** Neural speech decoding, neural speech prosthesis, Electrocorticographic (ECoG), speech synthesis

## Abstract

Decoding human speech from neural signals is essential for brain-computer interface (BCI) technologies restoring speech function in populations with neurological deficits. However, it remains a highly challenging task, compounded by the scarce availability of neural signals with corresponding speech, data complexity, and high dimensionality, and the limited publicly available source code. Here, we present a novel deep learning-based neural speech decoding framework that includes an ECoG Decoder that translates electrocorticographic (ECoG) signals from the cortex into interpretable speech parameters and a novel differentiable Speech Synthesizer that maps speech parameters to spectrograms. We develop a companion audio-to-audio auto-encoder consisting of a Speech Encoder and the same Speech Synthesizer to generate reference speech parameters to facilitate the ECoG Decoder training. This framework generates natural-sounding speech and is highly reproducible across a cohort of 48 participants. Among three neural network architectures for the ECoG Decoder, the 3D ResNet model has the best decoding performance (PCC=0.804) in predicting the original speech spectrogram, closely followed by the SWIN model (PCC=0.796). Our experimental results show that our models can decode speech with high correlation even when limited to only causal operations, which is necessary for adoption by real-time neural prostheses. We successfully decode speech in participants with either left or right hemisphere coverage, which could lead to speech prostheses in patients with speech deficits resulting from left hemisphere damage. Further, we use an occlusion analysis to identify cortical regions contributing to speech decoding across our models. Finally, we provide open-source code for our two-stage training pipeline along with associated preprocessing and visualization tools to enable reproducible research and drive research across the speech science and prostheses communities.

## Introduction

Speech loss due to neurological deficits is a severe disability that limits social and work life. Advances in machine learning and Brain-computer interface (BCI) systems have pushed the envelope to develop neural speech prostheses to enable people with speech loss to communicate[1–5]. An effective modality for acquiring data to develop such decoders involves Electrocorticographic (ECoG) recordings obtained in epilepsy surgery patients [4–10]. Implanted electrodes in Epilepsy patients provides a rare opportunity to collect cortical data during speech with high spatial and temporal resolution. Such approaches have revealed promising results in speech decoding [4, 5, 8–11].

Two challenges are inherent to speech decoding from neural signals. First, data to train neural-to-speech decoding are limited in duration, while deep learning models require extensive training data. Second, speech production varies in rate, intonation, pitch, etc., even within a single speaker producing the same word, complicating the underlying model representation [12, 13]. These challenges have led to diverse approaches to speech decoding with limited consistency in model approaches and the availability of public code to test and replicate findings across research groups.

Earlier approaches to decode and synthesize speech spectrograms from neural signals focused on linear models. These approaches achieved ∼ 0.6 Pearson correlation coefficient (PCC) or lower while providing simple model architectures that are easy to interpret and do not require large training datasets[14–16]. Recent research has focused on deep neural networks leveraging convolutional [8, 9] and recurrent [5, 10, 17] network architectures. These approaches vary across two major dimensions, the intermediate latent representation used to model speech and the quality of speech produced after synthesis. For example, cortical activity has been decoded into an articulatory movement space, which is then transformed into speech providing robust decoding performance but with a non-natural synthetic voice reconstruction [17]. Conversely, some approaches produced naturalistic reconstruction leveraging wavenet vocoders [8], General Adversarial Networks [11], and unit selection [18], but with limited accuracy. Further, most previous approaches employ non-causal architectures, which could limit real-time applications that inherently require causal operations. Finally, we are unaware of prior work providing an open-source code to facilitate the replication and comparison of decoding approaches.

To address these issues, we provide a novel ECoG-to-speech framework with a low-dimensional intermediate representation guided by participant-specific pre-training using speech signal only (Fig. 1). Our framework consists of an ECoG Decoder that maps the ECoG signals to interpretable acoustic speech parameters (e.g., pitch, voicing, and formant frequencies) and a Speech Synthesizer that translates the speech parameters to a spectrogram. The Speech Synthesizer is differentiable, enabling us to minimize the spectrogram reconstruction error during the training of the ECoG Decoder. The low-dimensional latent space, together with the guidance on the latent representation generated by a pre-trained Speech Encoder, overcomes data scarcity issues. Our publicly available framework produces naturalistic speech, and the ECoG decoder can be realized with different deep-learning model architectures and using different causality directions. Here we report this framework with multiple deep architectures (i.e., convolutional, recurrent, transformer) as the ECoG decoder and apply it to 48 neurosurgical patients. Our framework performs with high accuracy across models, with the best performance obtained by the convolutional (ResNet) architecture (Pearson Correlation of 0.804 between the original and decoded spectrogram). Our framework can achieve high accuracy using only causal processing and relatively low spatial sampling on the cortex. Further, we show comparable speech decoding from the grid implants on the left and right hemispheres, providing a proof of concept for neural prosthetics in patients suffering from expressive Aphasia (damage limited to the left hemisphere). Taken together, we provide a publicly available neural decoding pipeline with associated tools (https://xc1490.github.io/nsd/) to push forward research across the speech science and prostheses communities.

**Fig. 1.**
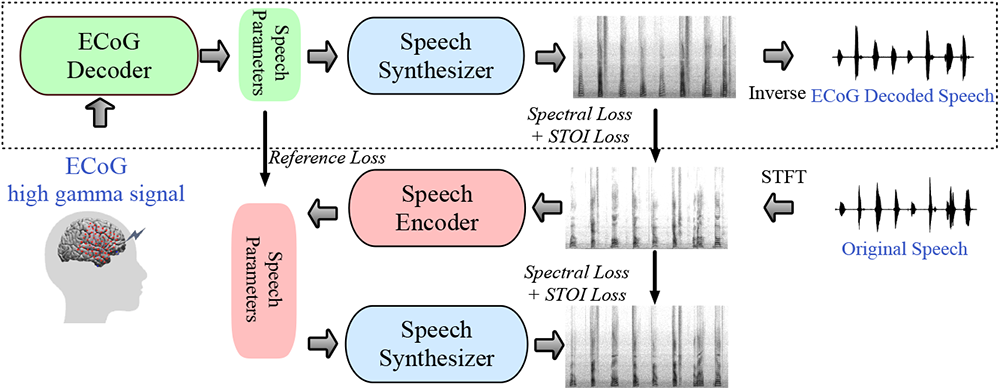
The proposed neural speech decoding framework. The upper part shows the ECoG-to-speech decoding pipeline. The ECoG Decoder generates time-varying speech parameters from ECoG signals. The Speech Synthesizer generates spectrograms from the speech parameters. A separate spectrogram inversion algorithm converts the spectrograms to speech waveforms. The lower part shows the speech-to-speech auto-encoder that generates the guidance for the speech parameters to be produced by the ECoG Decoder during the ECoG Decoder training. The Speech Encoder maps an input spectrogram to speech parameters, which are then fed to the same Speech Synthesizer to reproduce the spectrogram. The Speech Encoder and a few learnable participant-specific parameters in the Speech Synthesizer are pre-trained using speech signals only. Only the upper part is needed for decoding the speech from ECoG signals once the pipeline is trained.

### ECoG to Speech Decoding Framework and Our Training Pipeline

Our ECoG-to-Speech framework consists of an ECoG Decoder and a Speech Synthesizer, shown in the upper part of Fig. 1. The neural signals are fed into an ECoG Decoder that generates speech parameters, followed by a Speech Synthesizer, which translates the parameters into spectrograms (which are then converted to waveform). The training of our framework takes two steps. We first use semi-supervised learning on the speech signals alone. An autoencoder, shown in the lower part of Fig. 1, is trained so that the Speech Encoder derives speech parameters from a given spectrogram, while the Speech Synthesizer (used here as the decoder) reproduces the spectrogram from the speech parameters. Our Speech Synthesizer is fully differentiable and generates speech through a weighted combination of voice and unvoice speech components from input time series of speech parameters, including pitch, formant frequencies, loudness, etc. The Speech Synthesizer has only a few participant-specific parameters that are learned as part of the auto-encoder training. (see Methods section: Speech Synthesizer for more details). Currently, our Speech Encoder and Speech Synthesizer are participant-specific and can be trained using any speech signal of the participant, not limited to those with corresponding ECoG signals.

In the next stage, we train the ECoG Decoder in a supervised manner based on spectrogram losses (spectral and Short-Time Objective Intelligibility (STOI) [8, 19]) as well as guidance for the speech parameters generated by the pre-trained Speech Encoder (i.e., reference loss between speech parameters). By limiting the number of speech parameters (18 at each time step, see Methods Section: Summary of Speech Parameters) and using the reference loss, the ECoG Decoder can be trained with limited corresponding ECoG and speech data. Further, because our Speech Synthesizer is differentiable, we can back-propagate the spectral loss (differences between the original and decoded spectrograms) to update the ECoG Decoder. Here, we provide multiple ECoG Decoder architectures to choose from, including 3D ResNet [20], 3D SWIN Transformer [21], and LSTM [22]. Importantly, unlike many methods in the literature, we employ ECoG Decoders that can operate in a causal manner, which is necessary for real-time speech generation from neural signals. Note that once the ECoG Decoder and Speech Synthesizer are trained, they can be used for ECoG to speech decoding without using the Speech Encoder.

## Results

### Data

We employed our speech decoding framework across N=48 participants who consented to complete a series of speech tasks (see Methods Section: Datasets and Experiments). The participants were undergoing treatment for refractory Epilepsy with implanted electrodes for their clinical care. During the hospital stay, we acquired synchronized neural and acoustic speech data. ECoG data were obtained from five participants with hybrid-density(HB) sampling (clinical-research grid) and 43 participants with low-density(LD) sampling (standard clinical grid), who took part in five speech tasks: Auditory Repetition (AR), Auditory Naming (AN), Sentence Completion (SC), Word Reading (WR), and Picture Naming (PN). These tasks were designed to elicit the same set of spoken words across tasks while varying the stimulus modality. We provided 50 repeated unique words (400 total trials per participant), all of which were analyzed locked to the onset of speech production. We trained a model for each participant using 80% of available data for this participant and evaluated the model on the remaining 20% of data (with the exception of the more stringent word-level cross-validation).

### Speech Decoding Performance and Causality

We first aimed to directly compare decoding performance across different architectures, including those that have been employed in the neural speech decoding literature (recurrent and convolutional) and transformer-based models. While any decoder architecture could be used for the ECoG Decoder in our framework employing the same Speech Encoder guidance and Speech Synthesizer, we focused on three representative models for convolution (ResNet), recurrent (LSTM), and transformer (SWIN). Our results show that ResNet outperforms the other models providing the highest Pearson correlation coefficient (PCC) across N=48 participants (mean PCC=0.804, 0.798 for noncausal, causal respectively) but closely followed by SWIN (mean PCC=0.793, 0.796 for non-causal, causal respectively) shown in Fig. 2**a**. We find the same conclusion when evaluating three models using STOI+ [23], shown in Supplementary Fig. S1**a**. The causality of machine learning models for speech production has important implications for BCI applications. A causal model only uses past and current neural signals to generate speech. In contrast, non-causal models use past, present, and future neural signals. Previous reports typically employed non-causal models [5, 8, 10, 17], that could potentially use neural signals related to auditory and speech feedback, unavailable in realtime applications. Optimally, only the causal direction should be employed. We, therefore, compared the performance of the same models with either noncausal or causal temporal operations. Fig. 2**a** compares the decoding results of our models’ causal and non-causal versions. The causal ResNet model (PCC=0.798) achieved performance comparable to that of the non-causal model (PCC=0.804) with no significant differences between the two (Wilcoxon signed-rank test *p* = 0.1022). The same was true for the causal SWIN model (PCC=0.796) and its non-causal (PCC=0.793) counterpart (Wilcoxon signed-rank test *p* = 0.2794). In contrast, the performance of the causal LSTM model (PCC=0.713) was significantly inferior to that of its non-causal (PCC=0.757) version (Wilcoxon signed-rank test *p* = 0.0007). Further, the LSTM model showed consistently lower performance compared with ResNet and SWIN. However, we did not find significant differences between causal ResNet and causal SWIN performance (Wilcoxon signed-rank test *p* = 0.1413). Since the ResNet and SWIN models had the highest performance and were on par with each other and their causal counterparts, we chose to focus further analyses on these causal models, which we believe are best suited for prosthetic applications.

To ensure our framework can generalize well to unseen words, we added a more stringent word level cross-validation wherein random (10 unique) words were held entirely out during training (including both pre-training of Speech Encoder and Speech Synthesizer and training of ECoG Decoder). This ensures that different trials from the same word could not appear in both the training and testing sets. Results shown in Fig. 2**b** demonstrate that performance on the held-out words is comparable to our standard trial-based held-out approach (i.e., Fig. 2**a**. “ResNet”). It is encouraging that the model can decode unseen validation words well, regardless of which words were held out during the training.

Next, we show the performance of the ResNet causal decoder on the level of single words across two representative participants (low-density grids). The decoded spectrograms accurately preserve the spectro-temporal structure of the original speech (Fig. 2**c,d**). We also compare the decoded speech parameters with the reference parameters. For each parameter, we calculate the PCC between the decoded time series and the reference sequence showing an average PCC of 0.834 (voice weight, Fig. 2**d**), 0.545 (loudness, Fig. 2**e**), 0.907 (pitch, *f*_0_, Fig. 2**f**), 0.827 (first formant *f*_1_, Fig. 2**f**), 0.895 (second formant *f*_2_, Fig. 2**f**). The accurate reconstruction of the speech parameters, especially the pitch, voice weight, and first two formants, is essential for accurate speech decoding and naturalistic reconstruction that mimics the participant’s voice. We also provide a non-causal version of Fig. 2 in Supplementary Fig. S2. The fact that both non-causal and causal models could yield reasonable decoding results is encouraging.

**Fig. 2.**
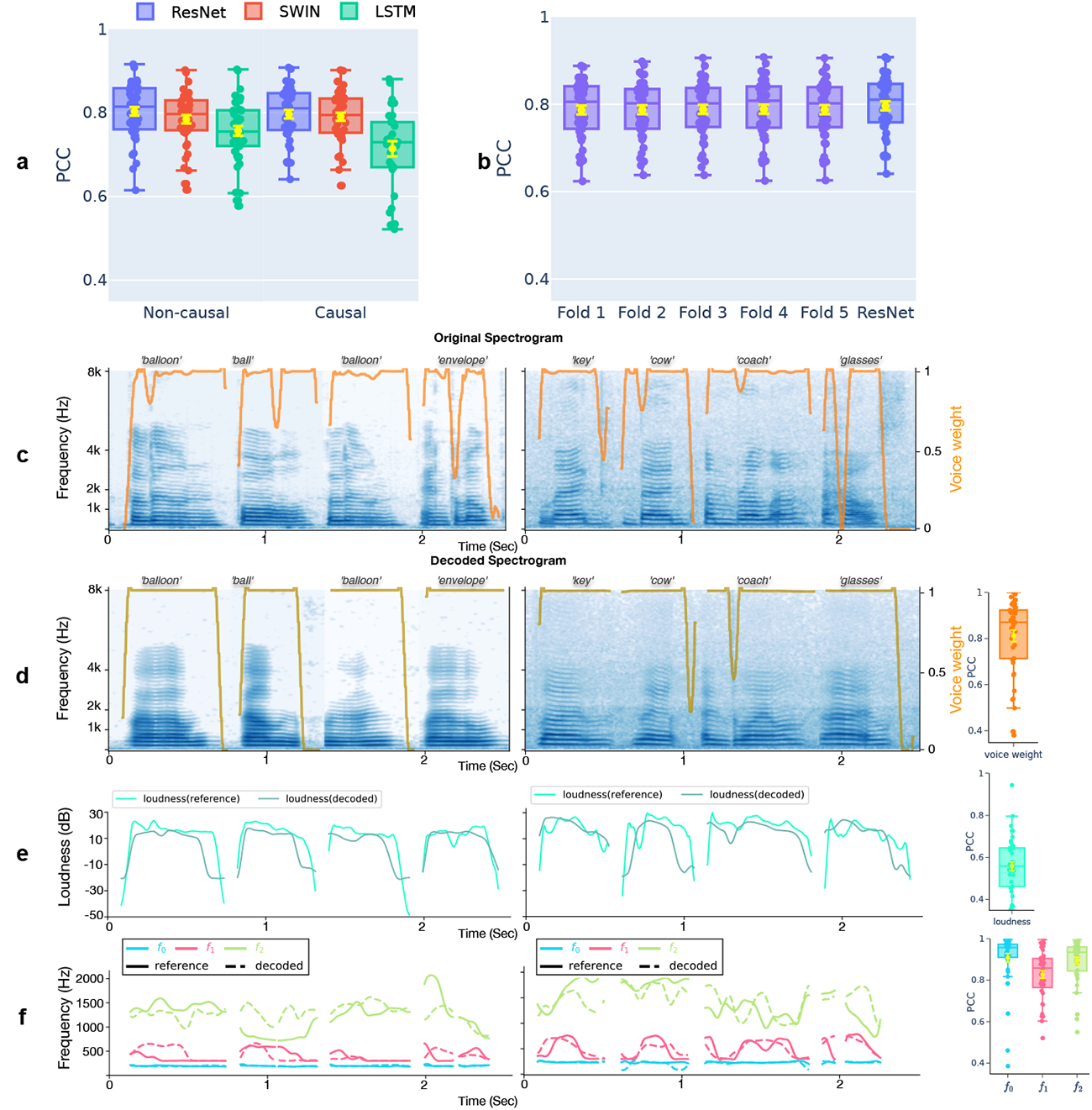
Decoding performance comparing the original and decoded spectrograms across non-causal and causal models. **a**, Performance of ResNet, SWIN, and LSTM models with non-causal and causal operations across all participants (n=48; 43 low-density ECoG grids and 5 hybrid density grids). The Pearson Correlation Coefficient (PCC) between the original and decoded spectrogram is evaluated on the held-out testing set and shown for each participant. Each data point in the box plot corresponds to a participant’s average PCC across all testing trials. **b,** A stringent cross-validation showing the Causal ResNet model performance on unseen words during training from 5-folds where we ensured the training and validation sets in each fold do not overlap in unique words. The performance across all five validation folds is comparable to our trial-based validation and denoted for comparison as ResNet (identical to the ResNet causal model in 2a). **c-f,** Example of decoded spectrograms and speech parameters by the causal ResNet model for eight words (from two participants) and the PCC values between the decoded and reference speech parameters across all participants. Spectrograms of the original (**c**) and decoded (**d**) speech are shown with orange curves overlaid representing the reference voice weight learned by the Speech Encoder (**c**) and the decoded voice weight by the ECoG Decoder (**d**). The correlations (PCC) between the decoded and reference Voice weights are shown on the right across all participants. (**e**) Decoded loudness parameter and reference loudness parameter are shown for the eight words as well as the PCC of the decoded loudness parameters across participants (boxplot on the right). (**f**) Decoded (dashed) and reference (solid) parameters for pitch (*f*_0_) and the first two formants (*f*_1_ and *f*_2_) are shown for the eight words as well as the PCC across participants (boxplots to the right). All boxplots represent the median, 25th and 75th quantiles across participants, and the yellow error bars denote the mean and standard error of the mean (across participants).

**Fig. 3.**
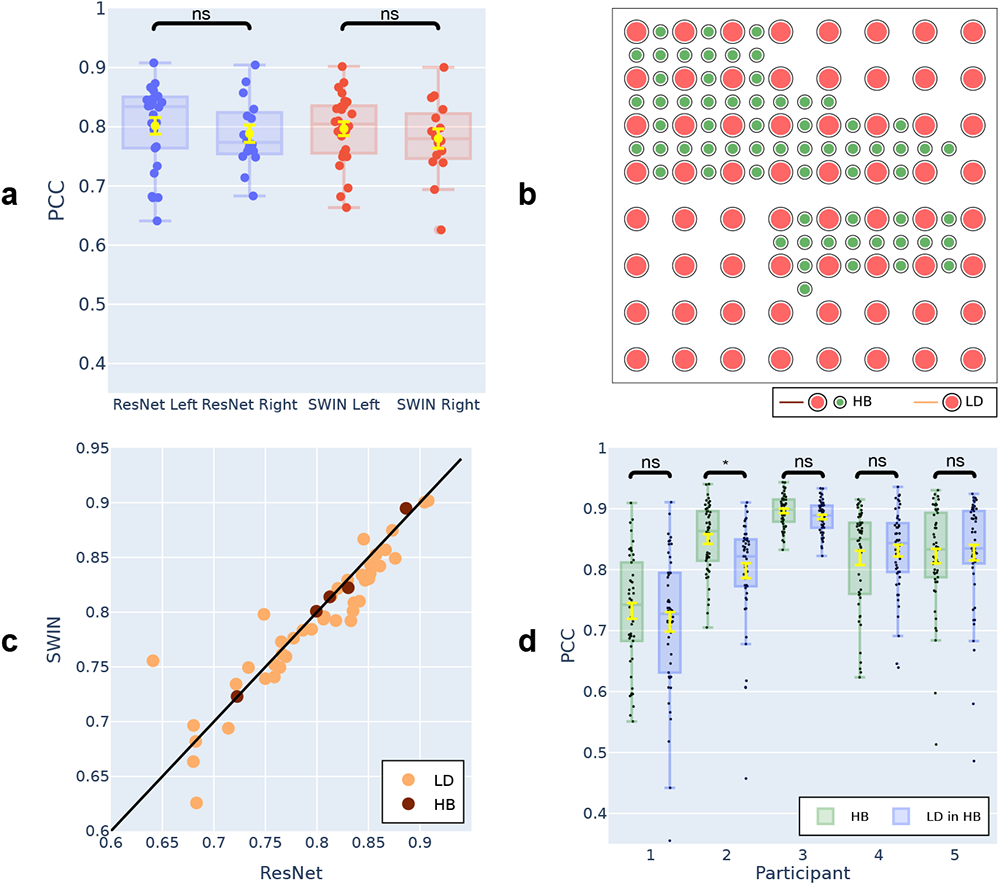
Comparison of decoding PCC under different settings of the 3D ResNet and 3D SWIN models. **a**, Comparison between left and right hemisphere participants, using causal models. No statistically significant differences in PCC exist between left (n=32) and right (n=16) hemisphere participants. **b,** An example hybrid density ECoG array, with a total of 128 electrodes. The 64 electrodes marked in red correspond to a low-density placement. The remaining 64 green electrodes, combined with red electrodes, reflect a hybrid density placement. **c,** Comparison between causal ResNet and causal SWIN models for the same participant across participants with hybrid-density (HB, n=5) or low-density (LD, n=43) ECoG grids. The two models have very similar decoding performances from the HB grids. The causal ResNet has slightly better performance on the LD grids. **d,** The decoding PCC by the ResNet model for HB participants when all electrodes are used vs when only LD-in-HB electrodes are considered. There are no statistically significant differences for 4 out of 5 participants.

### Left vs Right Hemisphere Decoding and Effect of Electrode Density

Most speech decoding studies focused on the language and speech-dominant left hemisphere ([24]). However, many patients who suffer damage to the left hemisphere are unable to speak ([25]), and right hemisphere decoding could serve as a valuable clinical target. To this end, we compared left vs right hemisphere decoding performance across our participants to establish the feasibility of a right hemisphere speech prosthetic. For both our ResNet and SWIN decoders, we found robust speech decoding from the right hemisphere (ResNet PCC=0.790, SWIN PCC=0.781), which were not significantly different from the left (Fig. 3**a**, ResNet Wilcoxon rank-sum test, p=0.312; SWIN Wilcoxon rank-sum test, p=0.325). A similar conclusion holds when evaluating STOI+ (Supplementary Fig. S1**b**, ResNet Wilcoxon rank-sum test, p=0.108; SWIN Wilcoxon rank-sum test, p=0.092). Next, we assessed the impact of the electrode sampling density for speech decoding, as many previous reports employ higher-density grids (0.4 mm) with more closely spaced contacts than typical clinical grids (1 cm). Five participants consented to hybrid grids (i.e., HB, see Fig. 3**b**), which had typical low density (i.e., LD) electrode sampling as well as additional electrodes interleaved. The hybrid density grids provided a decoding performance similar to clinical low-density grids in terms of PCC (Fig. 3**c**), with a slight advantage in STOI+, shown in Supplementary Fig. S3**b**. To ascertain if the additional spatial sampling indeed provides improved speech decoding, we compared models that decode speech based on all the hybrid electrodes vs. only the low-density electrodes in patients with HB grids (comparable to our other LD participants). Our findings (Fig. 3**d**) suggest that decoding results were not significantly different from each other (with the exception of Participant 2), including our evaluation of STOI+ (Fig. S3**c**). Together, these results suggest that our models can learn speech representations well from both the high and low spatial sampling of the cortex with the exciting finding of robust speech decoding from the right hemisphere.

**Fig. 4.**
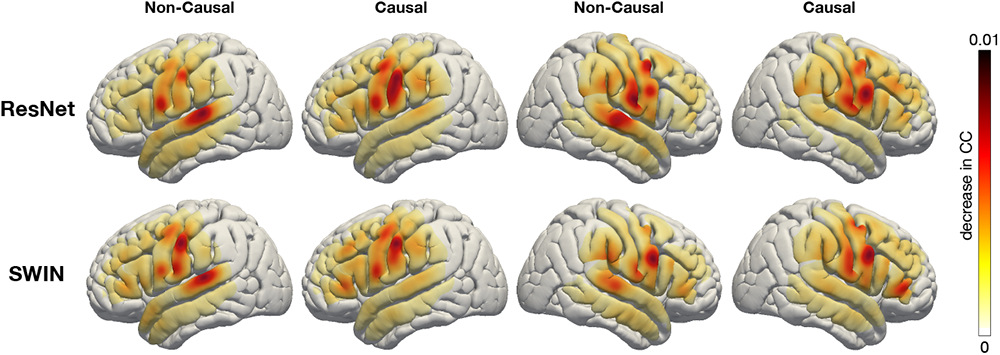
We visualize the contribution of each cortical location to the decoding result by both causal or non-causal decoding models through an occlusion analysis. The contribution of each electrode region in each participant is projected onto the standardized Montreal Neurological Institute (MNI) brain anatomical map and then averaged over all participants. Each subplot shows the causal or non-causal contribution of different cortical locations (red indicates higher contribution while yellow indicates lower contribution). For visualization purposes, we normalize the contribution of each electrode location by the local grid density since we have multiple participants with non-uniform density.

### Contribution analysis

Lastly, we investigate what cortical regions contribute to decoding, providing insight for targeted implantation in future prosthetics, especially on the right hemisphere, which has not been studied. We employed an occlusion approach to quantify the contributions of different cortical sites to speech decoding. If a region is involved in decoding, occluding the neural signal in the corresponding electrode (i.e., setting the signal to zero) would reduce the accuracy (PCC) of the reconstructed speech on the testing data (see Methods Section: Contribution Analysis). Therefore, we measure each region’s contribution by decoding PCC reduction when the corresponding electrode is occluded. We conducted this analysis across all electrodes and participants for causal and non-causal ResNet and SWIN decoders versions. The results shown in Fig. 4 show similar contribution values for both ResNet and SWIN models. The Non-Causal models show enhanced auditory cortex contributions compared with the Causal models, implicating auditory feedback in decoding and underlying the importance of employing only Causal models during speech decoding because neural feedback signals are not available for real-time decoding applications. Further, across the Causal models, both the right and left hemispheres show similar contributions across the sensorimotor cortex, especially on the ventral portion, which provides encouraging feasibility for right hemisphere neural prosthetics.

## Discussion

Our novel pipeline can decode speech from neural signals leveraging interchangeable architectures for the ECoG Decoder and a novel differentiable Speech Synthesizer. Our training process relies on estimating guidance speech parameters from the participant’s speech using a pre-trained Speech Encoder. This strategy enabled us to train ECoG Decoders with limited corresponding speech and neural data, which, when paired with our Speech Synthesizer, can produce natural-sounding speech. Our approach was highly reproducible across participants (N=48), providing evidence for successful causal decoding with convolutional (ResNet) and transformer (SWIN) architectures, both out-performing recurrent architecture (LSTM). Our framework can successfully decode from both high and low spatial sampling with high levels of decoding performance. Lastly, we provide one of the only reports of the robust speech decoding from the right hemisphere and the spatial contribution of cortical structures to decoding across the hemispheres.

Our decoding pipeline showed robust speech decoding across participants leading to a Pearson correlation coefficient within the range of 0.52-0.91 (Fig. 2**a**, Causal ResNet mean 0.798, median 0.81, range 0.64-0.91) between decoded and the ground truth speech across several architectures. We attribute our stable training and accurate decoding to the carefully designed components of our pipeline (e.g., Speech Synthesizer, Speech Parameter Guidance) and multiple improvements (see Methods Section: Speech Synthesizer, ECoG Decoder, and Model Training) over our previous approach on the subset of participants with hybrid density grids [26]. Previous reports investigated speech or text decoding used linear models [14, 15, 27], transitional probability [4, 28], recurrent neural networks [5, 10, 17], convolutional neural networks [8, 26], and other hybrid or selection approaches [9, 16, 18, 29, 30]. Overall, our results are similar to (or better than) most previous reports (48% of our participants showed 0.80-0.90 decoding PCC, see Fig. 3**c**). However, a direct comparison is complicated by multiple factors. Previous reports vary on reported performance metrics, as well as the stimuli decoded (e.g., continuous speech vs. single words) and the cortical sampling (i.e., high vs. low density, depth electrodes compared with surface grids). Our freely available pipeline, which can be used across multiple neural network architectures and tested on various performance metrics, can facilitate the research community to conduct more direct comparisons while still adhering to the highest standard of speech decoding.

The temporal causality of decoding operations, critical for real-time BCI applications, has not been considered by most prior studies. Many of these non-causal models relied on auditory (and somatosensory) feedback signals. Our analyses show that non-causal models rely on a robust contribution from STG, which is mostly eliminated by using a causal model (Fig. 4). We believe that non-causal models would show limited generalizability for real-time BCI applications due to their over-reliance on feedback signals that may be absent (if no delay is allowed) or incorrect (if a short latency is allowed during real-time decoding). Some approaches used imagined speech which avoids feedback during training [16], or showed generalizability to mimed production lacking auditory feedback [17]. However, most reports still employ non-causal models, which cannot rule out feedback during training. Indeed, our contribution maps show robust auditory cortex recruitment for non-causal Resnet and SWIN models (Fig. 4), in contrast to the causal counterparts, which decode based on more frontal regions. Further, recurrent neural networks widely used in the literature [5] are typically bi-directional, producing non-causal behaviors and longer latencies for prediction during real-time application. The recurrent network we tested performed the worst when trained with one direction (Causal LSTM, Fig. 2a). Our data suggest that causal convolutional and transformer models can perform on par with their non-causal counterparts and recruit more relevant cortical structures for decoding.

In our study, we leveraged an intermediate speech parameter space together with a novel differentiable Speech Synthesizer to decode participant-specific naturalistic speech (Fig. 1). Previous reports used varying approaches to model speech, including an intermediate kinematic space [17], an intermediate random vector (i.e., GAN) [11], or direct spectrogram representations [8, 17]. Our choice of speech parameters as the intermediate representation allowed us to represent participant-specific acoustics that drive the synthesis of a spectrogram as part of the end-end decoding architecture. Our Speech Synthesizer is motivated by popular vocoder models (generating speech by passing an excitation source through a filter [31] or sinusoidal [32]) and is fully differentiable, facilitating the training of the ECoG decoder using spectral losses through backpropagation. Unlike kinematic representations, this direct acoustic space produces natural-sounding speech that preserves patient-specific characteristics, which would be lost with the kinematic representation. Further, the guidance speech parameters can be obtained using a pretrained Speech Encoder that does not require neural data. Thus, it could feasibly be trained using older audio recordings or a proxy speaker model of choice in the case of patients without the ability to speak. Additionally, the low-dimensional acoustic space and pre-trained guidance using speech signals circumvent the data bottleneck in training ECoG to speech decoder and provide a highly interpretable latent space. Finally, our entire decoding pipeline is generalizable to unseen words (Fig. 2b). This provides an advantage compared to pattern matching approaches [18] that produce participant-specific utterances but with limited generalizability.

Many prior works employed high-density electrode coverage over the cortex, providing many distinct neural signals [5, 10, 17, 27]. One question we directly addressed was whether higher-density coverage improves decoding. Surprisingly, we found a high decoding performance in terms of spectrogram PCC for both low-density and higher (hybrid) density grid coverage (Fig. 3**c**). Further, comparing the decoding performance obtained using all electrodes in our hybrid-density participants vs. using only the low-density electrodes in the same participants showed that decoding did not significantly differ (Fig. 3**d**, albeit for one participant). We attribute these results to our ECoG decoder’s ability to learn speech parameters as long as there is sufficient peri-sylvian coverage, even in low-density participants.

A striking result was the robust decoding from right hemisphere cortical structures as well as the clear contribution of the right peri-sylvian cortex. Our results are consistent with the idea that syllable-level speech information is represented bilaterally [33]. However, our findings suggest that higher-level speech is well-represented in the right hemisphere. Our decoding results could directly lead to speech prostheses for patients who suffer from expressive aphasia or apraxia of speech. While our decoding provides evidence for a robust representation of speech in the right hemisphere, it is important to note that these regions may not be critical for speech. A minority of studies that have mapped both hemispheres for speech using electrical stimulation mapping found left dominance [34, 35]. Further, it remains unclear to what degree right hemisphere reorganization may affect decoding. While some studies have shown limited right hemisphere decoding of vowels ([36]) and sentences [37] the results were mostly mixed with left hemisphere signals. In contrast to decoding, some previous approaches have attempted non-invasive [38] as well as invasive [39] epidural stimulation to treat speech aphasia, which could be feasibly combined with a future neural prosthetic device. A minimally invasive decoding device is quite feasible, as recently demonstrated by gait decoding in bilateral motor cortices with epidural recordings that provided extraordinary results [40].

There are several limitations in our study. First, our decoding pipeline requires speech training data paired with ECoG recordings, which may not exist for paralyzed patients. This could be potentially mitigated provided the patient has older speech recordings (or by using a proxy speaker chosen by the patient) and neural recordings during imagined speech are used instead of overt speech. Second, our ECoG Decoder models (3D ResNet and 3D SWIN) assume a grid-based electrode sampling which may not be the case. Future work should develop model architectures that are capable of handling non-grid data, such as strips and depth electrodes (stereo intracranial EEG). Importantly such decoders could simply replace our current grid-based ECoG Decoders while still being trained using our overall pipeline. Lastly, we focused on word-level decoding, which may not be directly comparable to sentence-level decoding results of other studies.

To summarize, our neural decoding approach, capable of decoding natural-sounding speech from 48 participants, provides the following major contributions. First, our proposed latent representation using explicit speech parameters together with a differentiable Speech Synthesizer enables interpretable and intelligible speech decoding. Second, we directly consider the causality of the ECoG Decoder, providing strong support for causal decoding, which is essential for real-time BCI applications. Third, our promising decoding results using low sampling density and right hemisphere electrodes shed light on future neural prosthetic devices in patients with damage to the left hemisphere. Last, but not the least, we have made our framework open to the community with documentation (https://xc1490.github.io/nsd/), and we trust that this open platform will help propel the field forward, supporting reproducible science.

## Methods

### Datasets and Experiments

We collected neural data from 48 native English-speaking participants (26 female, 22 male) with refractory epilepsy who had electrocorticographic (ECoG) subdural electrode grids implanted at NYU Langone Hospital. Five participants had hybrid-density (HB) sampling, and 43 had low-density (LD) sampling. The ECoG array was implanted on the left hemisphere for 32 participants and on the right for 16. The Institutional Review Board of NYU Grossman School of Medicine approved all experimental procedures. After consulting with the clinical care provider, a research team member obtained written and oral consent from each participant. Each participant performed five tasks to produce target words in response to auditory or visual stimuli. The tasks were: Auditory Repetition (AR, repeating auditory words), Auditory Naming (AN, naming a word based on an auditory definition), Sentence Completion (SC, completing the last word of an auditory sentence), Visual Reading (VR, reading aloud written words), and Picture Naming (PN, naming a word based on a color drawing).

For each task, we used the exact 50 target words with different stimulus modalities (auditory, visual, etc.). Each word appeared once in the AN and SC tasks and twice in the others. The five tasks involved 400 trials with corresponding word production and ECoG recording for each participant. The average duration of the produced speech in each trial was 500 ms.

### Data collection and preprocessing

The study recorded ECoG signals from the perisylvian cortex (including STG, IFG, pre-central, and postcentral gyri) of 48 participants while they performed five speech tasks. A microphone recorded subjects’ speech and was synchronized to the clinical Neuroworks Quantum Amplifier (Natus Biomedical, Appleton, WI), which captured ECoG signals. The ECoG array consisted of 64 standard 8×8 macro contacts (10 mm spacing) for 43 participants with low-density sampling. For five participants with hybrid-density sampling, the ECoG array also included 64 additional interspersed smaller electrodes (1 mm) between the macro contacts (providing 10 mm center-to-center spacing between macro contacts and 5 mm center-to-center spacing between micro/macro contacts; PMT corporation, Chanassen, MN). See Fig. 3b. This FDA-approved array was manufactured for this study. A research team member informed participants that the additional contacts were for research purposes during consent. Clinical care solely determined placement location across participants (32 left hemispheres; 16 right hemispheres). The decoding models were trained separately for each participant using all trials except 10 randomly selected ones from each task, leading to 350 trials for training and 50 trials for testing. The reported results are for testing data only.

We sampled ECoG signals from each electrode at 2048 Hz and downsampled them to 512 Hz before processing. The electrodes with artifacts (e.g., line noise, poor contact with the cortex, high amplitude shifts) were rejected. The electrodes with interictal and epileptiform activity were also excluded from the analysis. A common average reference (across all remaining valid electrodes and time) was subtracted from each individual electrode. The electrodes with interictal and epileptiform activity were excluded from the analysis. After the subtraction, the Hilbert transform extracted the envelope of the high gamma (70-150 Hz) component from the raw signal, which is then down-sampled to 125 Hz. The signal of each electrode over a silent baseline of 250 ms before the stimulus served as the reference signal. Each electrode’s signal was normalized to the reference mean and variance (i.e., z-score). The signals of electrodes with artifacts or with interictal/epileptiform activity were set to 0. For participants with noisy speech recordings, we applied spectral gating to remove stationary noise from the speech using an open-source tool [41].

To pre-train the auto-encoder including the Speech Encoder and Speech Synthesizer, unlike our prior work in [26], which completely relied on unsupervised training, we provide supervision for some speech parameters to improve their estimation accuracy further. Specifically, we use the Praat method [42] to estimate pitch and four formant frequencies 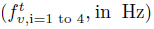 from the speech waveform. The estimated pitch and formant frequency are resampled to 125 Hz, the same as the ECoG signal and spectrogram sampling frequency. The mean square error between these speech parameters generated by the Speech Encoder and the ones estimated by the Praat method is used as a supervised reference loss, in addition to the unsupervised spectrogram reconstruction and STOI losses, making the training of the auto-encoder semi-supervised.

### Speech Synthesizer

Our Speech Synthesizer is inspired by the traditional speech vocoder, which generates speech by switching between voice and unvoice content, each generated by filtering a specific excitation signal. Instead of switching between the two components, we use a soft mix of the two components, making the Speech Synthesizer differentiable. This enables us to train the ECoG Decoder and the Speech Encoder end-to-end by minimizing the spectrogram reconstruction loss with back-propagation. Our Speech Synthesizer can generate a spectrogram from a compact set of speech parameters, enabling the training of the ECoG Decoder with limited data. As shown in Fig. 5, the synthesizer takes dynamic speech parameters as input and contains two pathways. The voice pathway applies a set of formant filters (each specified by center frequency 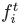, band-width 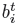 and amplitude 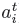) to the harmonic excitation (with pitch frequency *f_0_*) and generates the voice component, *V^t^*(*f*), for each time step *t* and frequency *f*. The noise pathway filters the input white noise with an unvoice filter (with center frequency 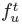, bandwidth 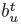 and amplitude 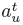) and produces the unvoice content, *U^t^*(*f*). The synthesizer combines the two components with a voice weight 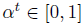 to obtain the combined spectrogram 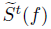 as follows:

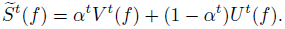

**Fig. 5.**
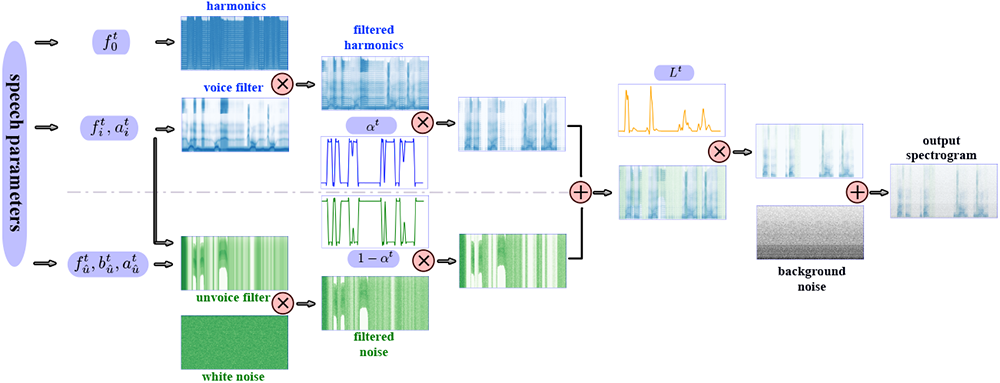
Differentiable Speech Synthesizer architecture. Our Speech synthesizer generates the spectrogram at time *t* by combining a voice component and an unvoice component based on a set of speech parameters at *t*. The upper part represents the voice pathway, which generates the voice component by passing a harmonic excitation with fundamental frequency 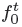 through a voice filter (which is the sum of 6 formant filters, each specified by formant frequency 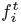 and amplitude 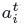). The lower part describes the noise pathway, which synthesizes the unvoice sound by passing white noise through an unvoice filter (defined by center frequency 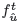 bandwidth 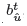 and amplitude 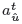. The two components are next mixed with voice weight *α^t^* and unvoice weight 1 *− α^t^*, respectively, and then amplified by loudness *L_t_*. A background noise (defined by a stationary spectrogram *B(f)*) is finally added to generate the output spectrogram. There are a total of 18 speech parameters at any time *t*, indicated in purple boxes.

The factor *α^t^* acts as a soft switch for the gradient to flow back through the synthesizer. The final speech spectrogram is

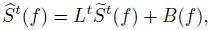

where *L^t^* is the loudness modulation and *B*(*f*) is the background noise. We describe the various components in more detail below.

### Formant filters in the voice pathway

We use multiple formant filters in the voice pathway to model formants that represent vowels and nasal information. The formant filters capture the resonance in the vocal tract, which can help recover a speaker’s timbre characteristics and generate natural-sounding speech. We assume the filter for each formant is time-varying and can be derived from a prototype filter *G_i_*(*f*), which achieves maximum at a center frequency 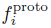 and has a half-power bandwidth 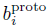. The prototype filters have learnable parameters and will be discussed later. The actual formant filter at any time is written as a shifted and scaled version of *G_i_*(*f*). Specifically, at time *t*, given an amplitude 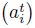, a center frequency 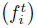, and a bandwidth 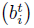, the frequency domain representation of the *i*-th formant filter is

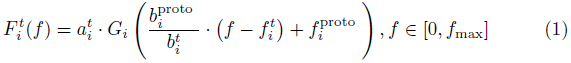

where *f*_max_ is half of the speech sampling frequency, which in our case is 8000 Hz.

Rather than letting the bandwidth parameters 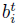 be independent variables, based on the empirically observed relationships between 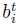 and the center frequencies 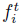, we set

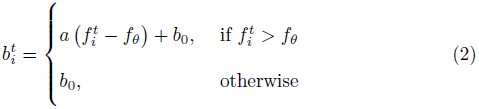

The threshold frequency *f_θ_*, slope *a*, and baseline bandwidth *b*_0_ are three parameters that are learned during the auto-encoder training, shared among all six formant filters. This parameterization helps to reduce the number of speech parameters to be estimated at every time sample, making the representation space more compact.

Finally the filter for the voice pathway with *N* formant filters is given by 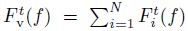. Previous studies have shown that two formants (*N* =2) are enough for intelligible reconstruction [43], but we use *N* =6 for more accurate synthesis in our experiments.

### Unvoice filters

We construct the unvoice filter by adding a single broadband filter 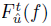 to the formant filters for each time step *t*. The broadband filter 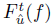 has the same form as equation (1) but has its own learned prototype filter 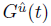 The speech parameters corresponding to the broadband filter include 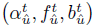. We do not impose a relationship between the center frequency 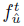 and the bandwidth 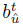. This allows more flexibility in shaping the broadband unvoice filter. But we constrain 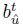 to be larger than 2000 Hz to capture the wide spectral range of obstruent phonemes. Instead of using only the broadband filter, we also retain the *N* formant filters in the voice pathway 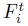 for the noise pathway. This is based on the observation that humans perceive consonants such as /p/ and /d/ not only by their initial bursts but also by their subsequent formant transitions until the next vowel [44]. We use identical formant filter parameters to encode these transitions. The overall unvoice filter is: 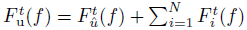.

### Voice Excitation

We use the voice filter in the voice pathway to modulate the harmonic excitation. Following [45], we define the harmonic excitation as 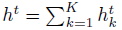, where *K*=80 is the number of harmonics.

The value of the *k*-th resonance at time step *t* is 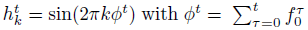, where 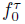 is the fundamental frequency at time *τ* . The spectrogram of *h^t^* forms the harmonic excitation in the frequency domain *H^t^*(*f*), and the voice excitation is 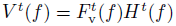.

### Noise Excitation

The noise pathway models consonant sounds (plosives and fricatives). It is generated by passing a stationary Gaussian white noise excitation through the unvoice filter. We first generate the noise signal *n*(*t*) in the time domain by sampling from the Gaussian process N (0, 1) and then obtain its spectrogram *N ^t^*(*f*). The spectrogram of the unvoice component is 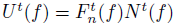.

### Summary of speech parameters and comparison with other Speech Synthesizers

The synthesizer generates the voice component at time *t* by driving a harmonic excitation with pitch frequency 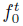 through *N* formant filters in voice pathway, each described by two parameters 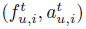. The unvoice component is generated by filtering a white noise through the unvoice filter consisting of an additional broadband filter with 3 parameters 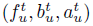. The two components are mixed based on the voice weight *α^t^* and further amplified by the loudness value *L^t^*. In total, the synthesizer input includes 18 speech parameters at each time step.

Unlike DDSP in [45], we do not directly assign amplitudes to the *K* harmonics. Instead, the amplitude in our model depends on the formant filters, which has two benefits:

- **The representation space is more compact.** DDSP requires 80 amplitude parameters 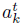 for each of the 80 harmonic components 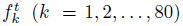 at each time step. In contrast, our synthesizer only needs a total of 18 parameters.
- **The representation is more disentangled.** For human speech, the vocal tract shape (affecting the formant filters) is largely independent of the vocal cord tension (which determines the pitch). Modeling these two separately leads to a disentangled representation.

In contrast, DDSP specifies the amplitude for each harmonic component directly resulting in entanglement and redundancy between these amplitudes. Furthermore, it remains uncertain whether the amplitudes 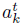 could be effectively controlled and encoded by the brain. In our approach, we explicitly model formant filters and fundamental frequency, which possess clear physical interpretations and are likely to be directly controlled by the brain. Our representation also enables a more robust and direct estimation of the pitch.

### Speaker-Specific Synthesizer Parameters

#### Prototype filters

Instead of using a pre-determined prototype formant filter shape, e.g., a standard Gaussian function, we learn a speaker-dependent prototype filter for each formant to allow more expressive and flexible formant filter shapes. We define the prototype filter *G_i_*(*f*) of the *i*-th formant as a piece-wise linear function, linearly interpolated from *g^i^*[*m*]*, m* = 1 *. . . M*, the amplitudes of the filter at *M* uniformly sampled frequencies in the range [0*, f*_max_]. We constrain g*^i^*[m] to increase and then decrease monotonically so that *G_i_*(*f*) is uni-modal and has a single peak value of 1. Given *g_i_*[*m*]*, m* = 1 *. . . M*, we can determine the peak frequency 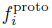 and the half-power bandwidth 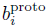 of *G_i_*(*f*).

The prototype parameters *g_i_*[*m*]*, m* = 1 *. . . M* of each formant filter are time-invariant and are determined during the auto-encoder training. Compared with [26], we increase *M* from 20 to 80 to enable more expressive formant filters, essential for synthesizing male speakers’ voices.

We similarly learn a prototype filter for the broadband filter *G_u_*(*f*) for the unvoice component, which is specified by *M* parameters *g_u_*(*m*).

#### Background Noise

The recorded sound typically contains background noise. We assume the background noise is stationary and has a specific frequency distribution, depending on the speech recording environment. This frequency distribution *B*(*f*) is described by *K* parameters, where *K* is the number of frequency bins (*K*=256 for females and 512 for males). The *K* parameters are also learned during the auto-encoder training. The background noise is added to the mixed speech components to generate the final speech spectrogram.

To summarize, our Speech Synthesizer has the following learnable parameters: the *M* = 80 prototype filter parameters for each of the *N* = 6 formant filters and the broadband filters (totaling *M* (*N* + 1) = 560), the three parameters *f_θ_, a, b*_0_ relating the center frequency and bandwidth for the formant filters (totaling 21), and *K* parameters for the background noise (256 for female and 512 for male). The total number of parameters for female speakers is 837, and that for male speakers is 1093. Note that these parameters are speaker-dependent but time-independent and can be learned together with the Speech Encoder during the training of the speech-to-speech auto-encoder, using the speaker’s speech only.

### Speech Encoder

The Speech Encoder extracts a set of (18) speech parameters at each time point from a given spectrogram, which are then fed to the speech synthesizer to reproduce the spectrogram.

We use a simple network architecture for the Speech Encoder, with temporal convolutional layers and multilayer perceptron (MLP) across channels at the same time point, as shown in Fig. 6**a**. We encode pitch *f_0_^t^* by combining features generated from linear and mel-scale spectrograms. The other 17 speech parameters are derived by applying temporal convolutional layers and channel MLP to the linear scale spectrogram. To generate formant filter center frequencies 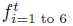, broadband filter frequency 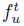, and pitch 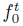, we use sigmoid activation at the end of the corresponding channel MLP to map the output to [0, 1], and then de-normalize it to real values by scaling [0, 1] to predefined [*f_min_, f_max_*]. The [*f_min_, f_max_*] values for each frequency parameter are chosen based on previous studies [46–49]. Our compact speech parameter space facilitates stable and easy training of our Speech Encoder.

**Fig. 6.**
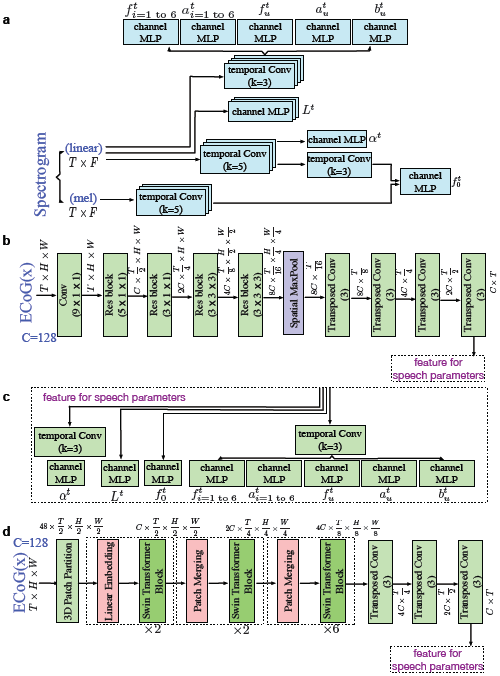
Speech Encoder and ECoG Decoder. **a,** Speech Encoder architecture. We input a spectrogram into a network of temporal convolution layers and channel MLPs that produce speech parameters. **b,** ECoG Decoder architecture using the 3D ResNet architecture. We first use several temporal and spatial convolutional layers with residual connections and spatiotemporal pooling to generate down-sampled latent features and then use corresponding transposed temporal convolutional layers to up-sample the features to the original temporal dimension. We then apply temporal convolution layers and channel MLPs to map the features to speech parameters, as shown in (c). **d,** ECoG Decoder using the 3D SWIN architecture. We use three or four stages of 3D SWIN blocks with spatial-temporal attention (3 blocks for LD and 4 blocks for HB) to extract the features from the ECoG signal. We then use the transposed versions of temporal convolution layers as in **b** to up-sample the features. The resulting features are mapped to the speech parameters using the same structure shown in (c).

### ECoG Decoder

We present the design details of two ECoG Decoders: the 3D ResNet ECoG Decoder and the 3D SWIN Transformer ECoG Decoder.

### 3D ResNet ECoG Decoder

This decoder adopts the ResNet architecture [20] for the feature extraction backbone of the decoder. Fig. 5**b** illustrates the feature extraction part. The model views the ECoG input as 3D tensors with spatiotemporal dimensions. In the first layer, we apply only temporal convolution to the signal from each electrode because the ECoG signal exhibits more temporal than spatial correlations. In the subsequent parts of the decoder, we have four residual blocks that extract spatiotemporal features using 3D convolution. After down-sampling the electrode dimension to 1 × 1 and the temporal dimension to *T/*16, we use several transposed Conv layers to up-sample the features to the original temporal size *T* . Fig. 5**c** shows how to generate the different speech parameters from the resulting features using different temporal convolution and channel MLP layers. The temporal convolution operation can be causal (i.e., using only past and current samples as input) or non-causal (i.e., using past, current, and future samples), leading to causal and non-causal models.

### 3D Swin Transformer ECoG Decoder

Swin Transformer[21] employs the window and shift window methods to enable self-attention of small patches within each window. This reduces the computational complexity and introduces the inductive bias of locality. Since our ECoG input data has three dimensions, we extend the Swin Transformer to three dimensions to enable local self-attention in both temporal and spatial dimensions among 3D patches. The local attention within each window gradually becomes global attention as the model merges neighboring patches in deeper Transformer stages.

Fig. 6**d** illustrates the overall architecture of the proposed 3D Swin Transformer. The input ECoG signal has a size of *T* ×*H*×*W*, where *T* is the number of frames and *H*×*W* is the number of electrodes at each frame. We treat each 3D patch of size 2×2×2 as a token in the 3D Swin Transformer. The 3D patch partitioning layer produces 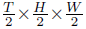 3D tokens, each with a *C* = 48 dimensional feature. A linear embedding layer then projects the features of each token to a higher dimension *C*(= 128).

The 3D Swin Transformer comprises three stages with 2, 2, 6 layers, respectively, for LD participants and four stages with 2, 2, 6, 2 layers for HB participants. It performs 2 × 2 × 2 spatial and temporal down-sampling in the patch merging layer of each stage. The patch merging layer concatenates the features of each group of 2 × 2 × 2 temporally and spatially adjacent tokens. It applies a linear layer to project the concatenated features to 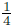 of their original dimension after merging. In the 3D Swin Transformer block, we replace the multi-head self-attention (MSA) module in the original Swin Transformer with the 3D shifted window multi-head self-attention module. It adapts the other components to 3D operations as well. A Swin Transformer block consists of a 3D-shifted window-based MSA module followed by a feed-forward network (FFN), a 2-layer MLP. Layer Normalization (LN) is applied before each MSA module and FFN, and a residual connection is applied after each module.

Consider a stage with *T* × *H* × *W* input tokens. If the 3D window size is P×M×M, we partition the input into 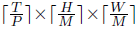 non-overlapping 3D windows evenly. We choose P = 16, M = 2. We perform the multi-head self-attention within each 3D window. However, this design lacks connection across adjacent windows, which may limit the representation power of the architecture. Therefore, we extend the shifted 2D window mechanism of the Swin Transformer to shifted 3D windows. In the second layer of the stage, we shift the window partition configuration by 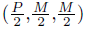 tokens along the temporal, height, and width axes from the previous layer. This creates cross-window connections for the self-attention module. This shifted 3D window design enables the interaction of electrodes with longer spatial and temporal distances by connecting neighboring tokens in non-overlapping 3D windows in the previous layer.

The temporal attention in the self-attention operation can be constrained to be causal (i.e., each token only attends to tokens temporally before it) or non-causal (i.e., each token can attend to tokens temporally before or after it), leading to the causal and non-causal models, respectively.

### Model Training

#### Training of the Speech Encoder and learnable parameters in the Speech Synthesizer

As described earlier, we pre-train the Speech Encoder and the learnable parameters in the Speech Synthesizer to perform a speech-to-speech auto-encoding task. We use multiple loss terms for the training. The modified multi-scale spectral loss is inspired by [45] and defined as

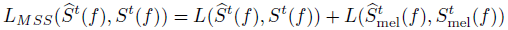

with

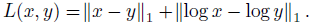

Here, *S^t^*(*f*) denotes the ground truth spectrogram and 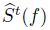 the reconstructed spectrogram in the linear scale, 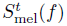 and 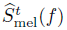 are the corresponding spectrograms in the mel-frequency scale. We sample the frequency range [0, 8000 Hz] with *K* = 256 bins for female participants. For male patients, we set *K* = 512 since they have lower *f*_0_ and it is better to have a higher resolution in frequency.

To improve the intelligibility of reconstructed speech, we also introduce the STOI loss by implementing the STOI+ metric [23], which is a variation of the original Short-Time Objective Intelligibility (STOI) metric [8, 19]. STOI+ [23] discards the normalization and clipping step in STOI and has been shown to perform best among intelligibility evaluation metrics. First, a one-third octave band analysis [19] is performed by grouping DFT bins into 15 one-third octave bands with the lowest center frequency set equal to 150 Hz and the highest center frequency equal to approximately 4.3 kHz. Let *x̂*(*k, m*) denote the *k^th^* DFT-bin of the *m^th^* frame of the ground truth speech. The norm of the *j*^th^ one-third octave band, referred to as a TF-unit, is then defined as:

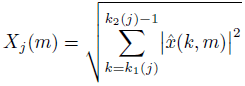

where *k*_1_(*j*) and *k*_2_(*j*) denote the one-third octave band edges rounded to the nearest DFT-bin. The TF representation of the processed speech *y*^ is obtained similarly and denoted by *Y_j_*(*m*). We then extract the short-time temporal envelopes in each band and frame, denoted *X_j,m_* and *Y_j,m_*, where 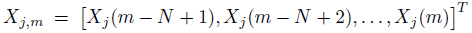, with *N* = 30. The STOI+ metric is the average of the Pearson correlation coefficient *d_j,m_* between *X_j,m_* and *Y_j,m_*, over all *j* and *m* [23]:

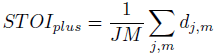

We use the negative of the STOI+ metric as the STOI loss:

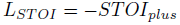

where *J* and *M* are the total numbers of frequency bins (J=15) and frames, respectively. Note that *L_ST OI_* is differentiable with respect to 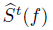, thus can be used to update the model parameters generating the predicted spectrogram 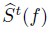.

To further improve the accuracy for estimating the pitch 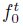 and formant frequencies 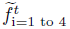, we add supervisions to them using the formant frequencies extracted by the Praat method [42]. The supervision loss is defined as

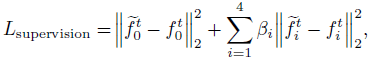

where the weights *β_i_* are chosen to be *β*_1_ = 0.1*, β*_2_ = 0.06*, β*_3_ = 0.03*, β*_4_ = 0.02, based on empirical trials. The overall training loss is defined as:

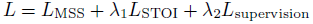

where weighting parameters *λ_i_* are empirically chosen to be *λ*_1_ = 1.2*, λ*_2_ = 0.1.

#### Training of the ECoG Decoder

With the reference speech parameters generated by the Speech Encoder and the target speech spectrograms as ground truth, the ECoG Decoder is trained to match these targets. Let us denote the decoded speech parameters as 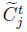, and their references as 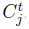, where *j* enumerates all speech parameters fed to the Speech Synthesizer. We define the reference loss as

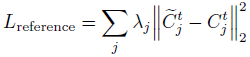

where weighting parameters *λ_j_* are chosen as follows: voice weight *λ_α_* = 1.8, loudness *λ_L_* = 1.5, pitch *λ_f_*_0_ = 0.4, formant frequencies *λ_f_*_1_ = 3*, λ_f_*_2_ = 1.8*, λ_f_*_3_ = 1.2*, λ_f_*_4_ = 0.9*, λ_f_*_5_ = 0.6*, λ_f_*_6_ = 0.3, formant amplitudes *λ_a_*_1_= 4*, λ_a_*_2_ = 2.4*, λ_a_*_3_= 1.2*, λ_a_*_4_= 0.9*, λ_a_*_5_= 0.6*, λ_a_*_6_= 0.3, broadband filter frequency *λ_fu_* = 10, amplitude *λ_au_*= 4, bandwidth *λ_bu_*= 4. Similar to speech-to-speech auto-encoding, we add supervision loss for pitch and formant frequencies derived by the Praat method and use the MSS and STOI loss to measure the difference between the reconstructed spectrograms and the ground truth spectrogram. The overall training loss for the ECoG Decoder is:

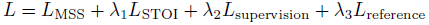

where weighting parameters *λ_i_* are manually chosen to be *λ*_1_ = 1.2*, λ*_2_ = 0.1*, λ*_3_ = 1.

We use the Adam optimizer [50] with hyper-parameters: *lr* = 10*^−^*^3^*, β*_1_ = 0.9*, β*_2_ = 0.999 to train both the auto-encoder (including the Speech Encoding and Speech Synthesizer) and the ECoG Decoder. We train a separate set of models for each participant. As mentioned earlier, we randomly select 50 out of 400 trials per participant as the test data and use the rest for training.

### Evaluation Metrics

In this paper, we use the Pearson Correlation Coefficient (PCC) between the decoded spectrogram and the actual speech spectrogram to evaluate the objective quality of the decoded speech, similar to [8, 18, 51].

We also use STOI+ [23] described in Method Section: Training of the ECoG Decoder to measure the intelligibility of decoded speech. The STOI+ value ranges from -1 to 1 and has been reported to have a monotonic relationship with speech intelligibility.

### Contribution Analysis Using the Occlusion Method

To measure the contribution of the cortex region under each electrode to the decoding performance, we adopt an occlusion-based method that calculates the change in the PCC between the decoded and the ground-truth spectrograms when an electrode signal is occluded (i.e., set to zeros), as in [26]. This method enables us to reveal the critical brain regions for speech production. We use the following notations: S*^t^*(*f*): the ground truth spectrogram; 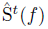: the decoded spectrogram with “intact” input (i.e., all ECoG signals are used); 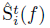: the decoded spectrogram with the *i*-th ECoG electrode signal being occluded; *r*(·, ·): correlation coefficient between two signals. The contribution of *i*-th electrode for a particular participant is defined as

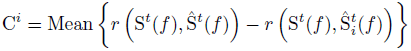

where Mean{·} denotes averaging across all testing trials of the participant.

We generate the contribution map on the standardized Montreal Neurological Institute (MNI) brain anatomical map by diffusing the contribution of each electrode of each participant (with a corresponding location in the MNI coordinate) into the adjacent area within the same anatomical region using a Gaussian kernel and then averaging the resulting map from all participants. To account for the non-uniform density of the electrodes in different regions and across the participants, we normalize the sum of the diffused contribution from all the electrodes at each brain location by the total number of electrodes in the region across all participants.

We estimate the noise level for the contribution map to assess the significance of our contribution analysis. To derive the noise level, we trained a shuffled model for each participant by randomly pairing the mismatched speech segment and ECoG segment in the training set. We derive the average contribution map from the shuffled models for all participants using the same occlusion analysis described earlier.

Contribution levels below the noise levels at corresponding cortex locations are assigned a value of 0 (white) in Fig. 4.

## Acknowledgments

This work was supported by the National Science Foundation under Grant No. IIS-1912286 (Y.W., A.F.) and National Institute of Health R01NS109367, R01NS115929, R01DC018805 (A.F.).

