## supplementary file for "A Neural Speech Decoding Framework Leveraging Deep Learning and Speech Synthesis"

The main manuscript presented the decoding performance in terms of the PCC. Here we present additional evaluations using the STOI+ metric, which has a better correlation with the intelligibility of decoded speech. We also present additional results obtained with the non-causal model.

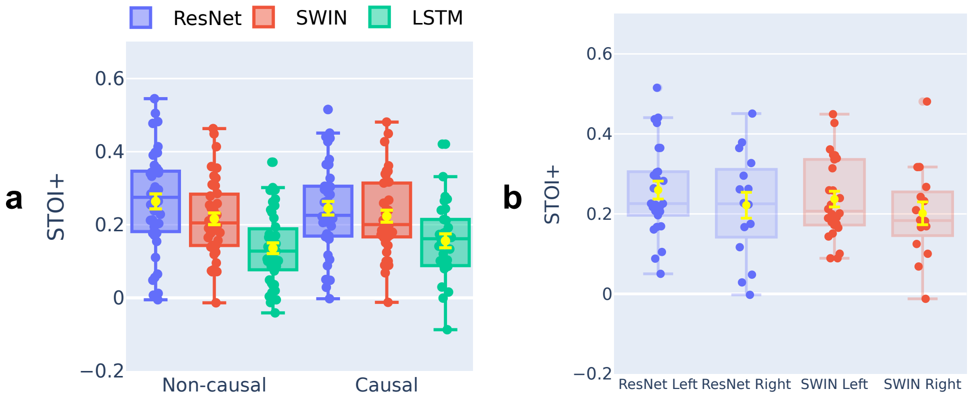


**Fig. S1.** | **Comparison of decoding STOI+ under different settings of the 3D ResNet, 3D SWIN, and LSTM models.** **a,** Performance of ResNet, SWIN, and LSTM models with non-causal and causal operations across all participants (n=48; 43 low-density ECoG grids and 5 hybrid density grids). The STOI+ between the original and decoded spectrogram is evaluated on the held-out testing set and shown for each participant. Each data point corresponds to a participant's average PCC across all testing trials. The boxplot represents the median, 25th and 75th quantiles across participants, and the yellow error bar denotes the mean and standard error of the mean. As with PCC (Fig. 2a in the manuscript), ResNet and SWIN models have similar performance, but the LSTM model is significantly worse. **b,** STOI+ comparison between left and right hemisphere participants, using causal ResNet and SWIN models. No statistically significant decoding performance differences exist between left (n=32) and right (n=16) hemisphere participants (ResNet Wilcoxon rank-sum test, p=0.108; SWIN Wilcoxon rank-sum test, p=0.092), although the left hemisphere participants have slightly greater mean STOI+.

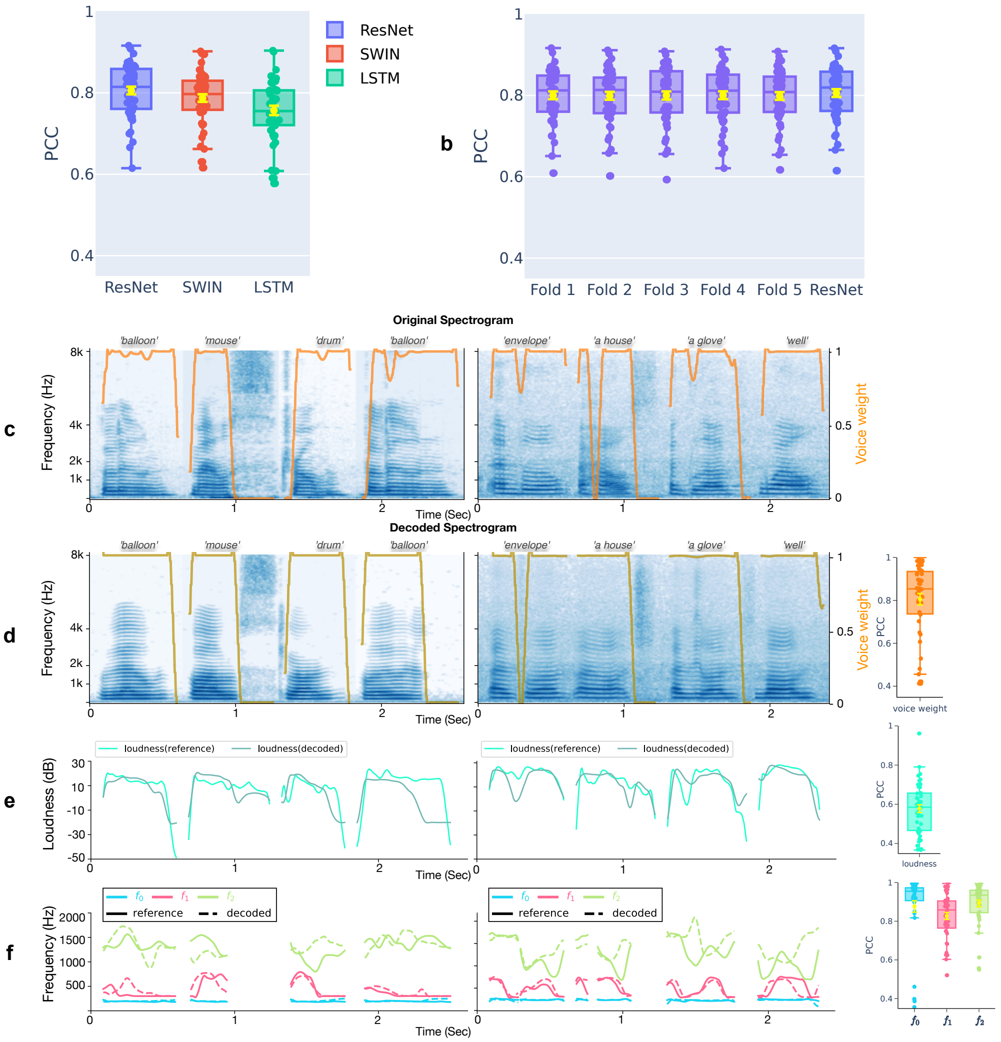


**Fig. S2.** | **Pearson Correlation Coefficient (PCC) between the original and decoded spectrograms by non-causal models. a,** PCC distributions of 3D ResNet, 3D SWIN, and LSTM models for all participants (n=48), including 43 participants with low-density ECoG grids and 5 participants with hybrid density grids, evaluated on the held-out testing set, which contains different speech trials from the training set but include overlapping words. **b,** ResNet model PCC on unseen words during training from a 5-fold cross-validation study where the training and validation sets in each fold have non-overlapping words. The performance across all five validation folds is similar to randomly selected test trials, demonstrating the model's generalization ability. **c-f,** Example decoded spectrograms and speech parameters by the non-causal ResNet model for four words each from two participants and PCC between the decoded and reference speech parameters across all participant trials. **c,d,** Comparison of original (**c**) and decoded (**d**) spectrograms. The orange curves overlaid on the spectrograms in **c** and **d** show the reference voice weight generated by the speech encoder and the decoded voice weight by the ECoG decoder, respectively. The box plot in **d** shows the PCC between the decoded and reference voice weight for all participants. **e,** Decoded loudness parameter (green) compared to reference loudness parameter (light green). The box plot shows the PCC of the decoded and reference loudness parameters. **f,** Comparison of the decoded (dashed) and reference (solid) pitch ($f_0$) and first two formants ($f_1, f_2$). The results achieved with non-causal models follow a very similar trend as those obtained with causal models, shown in Fig. 2f.

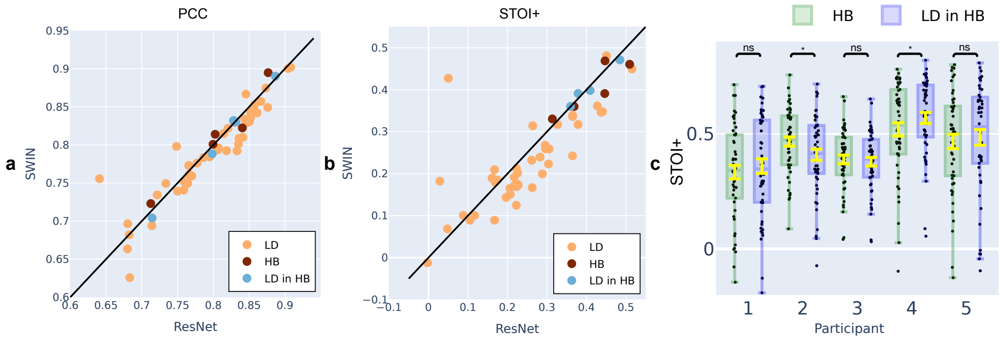


**Fig. S3.** | **Comparison of decoding PCC and STOI+ of the causal 3D ResNet and 3D SWIN models for the same participant across participants with hybrid-density (HB, n=5), low-density (LD, n=43), and only LD-in-HB ECoG grids.** In terms of PCC (**a**), both models have similar decoding performances on both HB, LD, and LD-in-HB participants. In terms of STOI+ (**b**), both models have slightly better performance on the HB participants, and the ResNet has better performance than the SWIN model for both HB and LD participants, but the difference is not statistically significant, as shown in Fig. S1a. The LD-in-HB and HB participants have very similar performances in both PCC and STOI+ (**c**). The decoding STOI+ by the ResNet model for HB participants when all electrodes are used vs when only LD-in-HB electrodes are considered. There are no statistically significant differences for 3 out of 5 participants. In 3 out of 5 participants, using LD-in-HB electrodes even gives us higher STOI+ compared with using HB only

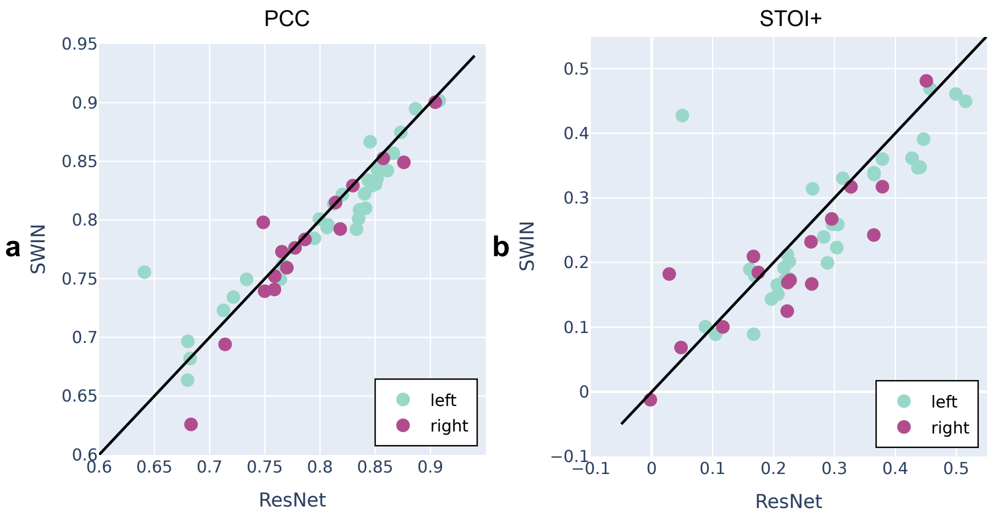


**Fig. S4.** | **Comparison of decoding PCC and STOI+ of the causal 3D ResNet and 3D SWIN models for the same participant across participants with right hemisphere (n=16) or left hemisphere (n=32) grids** (**a**) In terms of PCC, both models have similar decoding performances on the left and right hemispheres (ResNet Wilcoxon rank-sum test, p=0.312; SWIN Wilcoxon rank-sum test, p=0.325). (**b**) In terms of STOI+, both models have slightly better performance for the left hemisphere decoding, and the ResNet model has better performance for both left and right hemisphere decoding, but the difference is not statistically significant, as shown in Fig. S1b. (ResNet Wilcoxon rank-sum test, p=0.108; SWIN Wilcoxon rank-sum test, p=0.092)
